# Distinct gut metagenomics and metaproteomics signatures in prediabetics and treatment-naïve type 2 diabetics

**DOI:** 10.1101/666263

**Authors:** Huanzi Zhong, Huahui Ren, Yan Lu, Chao Fang, Guixue Hou, Ziyi Yang, Bing Chen, Fangming Yang, Yue Zhao, Zhun Shi, Baojin Zhou, Jiegen Wu, Hua Zou, Jin Zi, Jiayu Chen, Xiao Bao, Yihe Hu, Yan Gao, Jun Zhang, Xun Xu, Yong Hou, Huanming Yang, Jian Wang, Siqi Liu, Huijue Jia, Lise Madsen, Susanne Brix, Fang Liu, Karsten Kristiansen, Junhua Li

## Abstract

**Background:** The gut microbiota plays important roles in modulating host metabolism. Previous studies have demonstrated differences in the gut microbiome of T2D and prediabetic individuals compared to healthy individuals, with distinct disease-related microbial profiles being reported in groups of different age and ethnicity. However, confounding factors such as anti-diabetic medication hamper identification of the gut microbial changes in disease development.

**Method:** We used a combination of in-depth metagenomics and metaproteomics analyses of faecal samples from treatment-naïve type 2 diabetic (TN-T2D, n=77), pre-diabetic (Pre-DM, n=80), and normal glucose tolerant (NGT, n=97) individuals to investigate compositional and functional changes of the gut microbiota and the faecal content of microbial and host proteins in Pre-DM and treatment-naïve T2D individuals to elucidate possible host-microbial interplays characterising different disease stages.

**Findings:** We observed distinct differences characterizing the gut microbiota of these three groups and validated several key features in an independent TN-T2D cohort. We also demonstrated that the content of several human antimicrobial peptides and pancreatic enzymes differed in faecal samples between three groups, such as reduced faecal level of antimicrobial peptides and pancreatic enzymes in TN-T2D.

**Interpretation:** Our findings suggest a complex, disease stage-dependent interplay between the gut microbiota and the host and emphasize the value of metaproteomics to gain further insight into interplays between the gut microbiota and the host.

**Funding:** National Key Research and Development Program of China, No. 2017YFC0909703, Shenzhen Municipal Government of China, No. JCYJ20170817145809215, and National Natural Science Foundation of China, No. 31601073.

## Introduction

Type 2 diabetes mellitus (T2D) is a chronic heterogeneous disorder associated with hyperglycaemia and low grade inflammation [1,2]. The prevalence has increased dramatically in Westernized countries, and also in China, where 11.6% and 36% of Chinese adults suffer from diabetes and prediabetes (Pre-DM), respectively [3]. Due to complications and comorbidities related to the development of T2D, comprehensive characterization of phenotypic, metabolic and molecular changes of the host and the gut microbiota in pre-DM and T2D compared to NGT is needed to enable early identification of prediabetic individuals at high risk of T2D development. Cross-sectional metagenomic studies have linked alterations in the gut microbiome to T2D and prediabetes [4–7]. However, a few recent intervention studies have reported profound impact of antidiabetic drugs on the human gut microbiome, such as metformin, acarbose and glucagon-like peptide-1 (GLP-1) based therapies [8–13], emphasizing the importance of controlling for medication in studies on association between the microbiota and T2D. Moreover, distinct disease-related microbial profiles have been reported in different age and ethnic groups [4–7], making it difficult to identify the microbes possibly involved in disease development. Thus, detailed information on the gut microbial species associated with T2D onset and progression is still limited. Whereas information from metagenomics is limited to identification of the presence of genes, taxa, and their inferred functional capacity, introduction of additional omics approaches including metabolomics, metatranscriptomics, and metaproteomics have increased our knowledge of microbial activity in health and disease [14–17]. For instance, recent metatranscriptomics studies on inflammatory bowel disease and cirrhosis cohorts have revealed considerable discrepancies between data obtained from metagenomics vs metatranscriptomics analyses [17,18]. As metaproteomics enables identification of microbial and human proteins simultaneously in faecal samples [14,19,20], such an approach offers a potential for deciphering both active microbial functions and host-microbiota interactions.

In the present study, we examined 254 stool samples collected from a Chinese cohort combining shotgun metagenomics and metaproteomics analyses. We characterized substantial differences between NGT, Pre-DM and TN-T2D individuals. Of note, consistent aberrations in Pre-DM and TN-T2D individuals included lower abundances of *Clostridiales* species and higher abundances of *Megasphaera elsdenii* compared to NGT individuals. Several robust microbial compositional changes were detected at both the DNA and protein levels, such as an enrichment of *E. coli* in Pre-DM individuals and an increased abundance of *Bacteroides spp*. in TN-T2D patients. Several Pre-DM-specific features were furthermore uncovered, including a reduced capacity for processes involved in energy metabolism and bacterial growth, and an enrichment of *Prevotella* proteins as detected by metaproteomics. Thus, our findings revealed distinct characteristics of the intestinal ecosystem in the Pre-DM stage. Of note, proteomics analyses of the faecal samples revealed that the levels of a number of human proteins including several antimicrobial peptides (AMPs) differed in faecal samples from NGT, Pre-DM, and TN-T2D individuals, suggesting that specific differences in the host response amongst groups might also influence the composition of the gut microbiota, or vice versa. In conclusion, our study provides a basis for further analyses integrating faecal metagenomics and metaproteomics which may lead to a better understanding of mechanisms underlying the development of Pre-DM and T2D.

## Materials and Methods

### Suzhou T2D study population

The study population recruited from community residents from Suzhou, comprised 97 Chinese adults with normal glucose tolerance (NGT), 80 prediabetes patients (Pre-DM) and 77 newly diagnosed, treatment naïve type 2 diabetes patients (TN-T2D). All TN-T2D patients and Pre-DM individuals were screened and newly diagnosed according to the 2011 WHO criteria via well-trained staffs from the Suzhou Centre for Disease Prevention and Control (CDC), as described in detail in a recent published lipidomic study based on this cohort [21]. All enrolled 254 individuals have reported with no anti-diabetic treatments; thus, none have had taken insulin, or any oral or injectable anti-diabetic medication before. Stool samples for metagenomics were self-collected in 2ml faecal containers and immediately stored at −80°C and transported to the laboratory on dry ice. DNA was extracted as previously described [4]. A summary of sample information is presented in **Table S1**. In addition, shotgun metagenomic datasets of stools from 94 anti-diabetic medication TN-T2D patients from Shanghai [9], a city near to Suzhou, were used for validation purpose.

### Method for Metagenomics

#### 1. Generation of BGISEQ-500 based faecal metagenome data set

In this study, we performed DNA library construction and the combinatorial probe-anchor synthesis (cPAS)-based BGISEQ-500 sequencing for metagenomics (single-end; read length of 100bp) and applied the same quality control workflow to filter the low-quality reads in accordance with the recently published metagenomic study using this new platform [22]. The remaining high-quality reads were then aligned to hg19 to remove human reads [23]. Metagenomic data statistics is provided in **Table S2**.

#### 2. Profiling of metagenomic samples and biodiversity analysis

High-quality non-human reads were aligned to the 9.9M integrated gene catalogue (IGC) by SOAP2 using the criterion of identity ≥ 90% [23]. Sequence-based gene abundance profiling was performed as previously described. The relative abundances of phyla, genera, species and KOs were calculated by the sum of the relative abundance of their annotated genes. The alpha diversity (within-sample diversity) was quantified by the Shannon index using the relative abundance profiles at gene, genus and KO levels as described [23]. The beta diversity (between-sample diversity) was calculated using Bray-Curtis dissimilarity (R version 3.3.2, vegan package 2.4-4).

#### 3. Metagenome-wide association analysis (MWAS)

MWAS was performed on the Suzhou T2D cohort as previously described [4]. Using non-parametric Kruskal-Wallis test (R version 3.3.2 stats package), we identified 266,015 genes showing significant different abundances between the NGT, Pre-DM and TN-T2D groups (*P* < 0.05). After clustering, a total of 126 MLGs (≥100 genes) were generated from these genes. The relative abundance of each MLG was summed using the relative abundance values of all genes from this MLG. The taxonomic annotation of each MLG was determined if more than 50% of genes in this MLG could be assigned to a certain taxon according to their IGC annotation. The genes of 85 unclassified MLGs were further annotated using a reference sequence database including 1520 high-quality genomes cultivated from healthy Chinese individuals [24], resulted in the taxonomic annotations of 11 additional MLGs (See detailed information in **Table S5**).

### Method for Metaproteomics

#### 1. Sample preparation and LC-MS/MS analysis

Faecal samples from 84 individuals from NGT, Pre-DM, and TN-T2D individuals were used for metaproteome analysis using isobaric tags for relative and absolute quantitation (iTRAQ)–coupled-liquid chromatography tandem mass spectrometry (LC-MS/MS) (**Figure S1**). Each group consisted of 28 randomly selected individual samples with matched age, sex and BMI by propensity score matching (R version 3.3.2, MatchIt package 2.4-21) [25] (**Table S3**). Faecal samples were processed using the filter-aided sample preparation (FASP) protocol [26]. Briefly, 100mg frozen faeces from each individual were suspended in 500μl lysis buffer (4% SDS, 100mM dithiothreitol, 100mM Tris-HCL (pH=7.8) with freshly added protease inhibitors (cOmplete™, EDTA-free Protease Inhibitor Cocktail, Roche Applied Science). The samples were incubated for 5 min at 100 °C, followed by sonication to decrease the viscosity. The protein supernatants were collected after centrifugation at 30,000g at 4 °C for 30 min and then quantified using a 2D-quant kit (Sigma). For each diagnostic group, protein extracts in equal amounts from four individuals were pooled, and the selected 28 samples were thus aliquoted into 7 mixtures. A reference sample was created by pooling equal amounts of protein from each of 84 individual sample and 28 samples from self-reported T2D patients. Each mixture containing 100μg proteins was loaded onto a 10 kDa cut-off spin column (Vivacon 500, Sartorius AG, Goettingen, Germany). The lysate was adjusted to 8M urea by centrifuging to remove SDS and low-molecular-weight material. After reduction by dithiothreitol (DTT) and alkylation by iodoacetamide (IAM), 8M urea was added and centrifuged to remove any remaining reagent such as IAM. The urea buffer was then replaced with 0.5M triethylammonium bicarbonate (TEAB) and the sample was washed with 0.5M TEAB 5 times. Trypsin (Promega, Madison, WI, USA) was added to digest the protein at a protein: trypsin ratio of 50:1 and the mixtures were incubated for 18 hours at 37 °C. The resulting peptides were eluted twice with 100μl 0.5M TEAB by centrifuging at 12,000 g for 30 min and vacuum-dried. The peptide mixture samples were then dissolved in 0.5M TEAB and labelled with 8-plex iTRAQ reagents according to the manufacturer’s protocol (AB Sciex, USA). For each diagnostic group, 7 mixtures were labelled with tags from I113 to I119. To perform the iTRAQ quantitation throughout the whole experiment, we labelled the reference sample by tag 121 in each iTRAQ run. Thus, three independent 8-plex iTRAQ runs were conducted. Subsequently, labelled peptides were separated on a LC-20AB HPLC system (Shimadzu, Kyoto, Japan) with an Ultremex SCX column (Phenomenon, Torrance, CA) and collected into 20 fractions. Each fraction was analysed via a NanoLC system coupled with a Q Exactive mass spectrometry (Thermo Fisher Scientific, San Jose, CA) as described previously [27].

#### 2. Database searching and protein identification

For protein database searching, we used Mascot (Version 2.3) [28] as the search engine with the following parameters: trypsin was used as default enzyme and up to two missed cleavages were allowed. Carbamidomethyl (C), iTRAQ8plex (N-term) and iTRAQ8plex (K) were chosen as fixed modifications, and Oxidation (M) was chosen as variable modification. The peptide mass tolerance was set to 10 ppm and the fragment mass tolerance to 0.03 Da.

A two-step search method was applied. The MS/MS spectra were first searched against a collection of three protein sequence databases, including *Homo sapiens* sequences retrieved from SwissProt (release 2014_11), and human gut microbial protein sequences of IGC genes mapped by sequencing reads from our 254 metagenomic samples. The detailed search parameters are presented in **Table S4**. The Mascot search yielded a set of scored peptide-spectrum matches (PSMs) and the proteins were inferred from the PSMs. Subsequently, a target-decoy protein database was created containing the above-mentioned proteins and the reversed sequences from these proteins. A second round search based on the target-decoy database was performed to control for false positives as described elsewhere [29]. The PSMs were re-scored by Mascot Percolator [30] integrated into IQuant [31], and filtered at false discovery rate (FDR) ≤ 0.01. To improve the confidence in identification, peptides supported by ≥ 2 spectra were retained and protein identifications were thus inferred.

#### 3. Meta-protein Generation

Due to the shared similarity of metagenomic protein reference sequences, a microbial peptide hit is typically returned from several proteins within and between species. To avoid inflating numbers and alleviate taxonomic ambiguities of identified microbial proteins, several processes were performed to reduce data redundancy. We first grouped the microbial proteins with at least one shared peptide to generate protein clusters (**Figure S2**). Each cluster was then processed according to the maximum parsimony principle. The minimum protein sets containing all peptides of each cluster were selected and defined as the meta-protein representing the cluster (**Figure S2**). Individual proteins which only contained unique peptides were also assigned as a meta-protein. All redundant non-meta-protein sequences were thus omitted in subsequent analyses.

#### 4. Protein Quantification

Protein quantification was performed by IQuant [31] in the following three steps. We first normalized the intensities of iTRAQ reporter ions for all spectra across the eight iTRAQ-labelled samples (I113…I119, I121) using the formula (1) as follows:

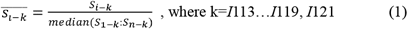

Where 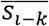 is the normalized relative intensity of spectrum *i* in the label *k*.

The reporter ion ratios were then determined using the formula (2):

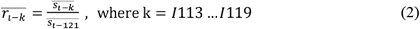

Where 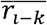 is the ratio of relative intensity of spectrum *i* in the label *k*, with *S*_*i*−121_, the relative intensity of the global QC labelled with 121 tags, as denominators.

For protein quantification, only unique peptides were taken into consideration. The relative protein ratio was calculated using the mean relative intensity ratio of all unique peptide spectra in each protein using the formula (3):

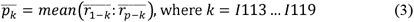

Where 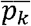 is the protein ratio in label K and acts as an indication of the relative proportions of that protein between the differently labelled samples.

#### 5. Protein annotation

For microbial meta-proteins, taxonomic and functional annotations of identified proteins were derived from the putative protein-coding IGC genes. As a result, we linked 64.15% (8777 of 11,980) of the meta-proteins with annotation at the phylum or lower taxonomical levels and 80.27% (10983 of 11,980) with KEGG Ontology (KO) annotation. For human proteins, functional annotations were obtained from UniProtKB/Swiss-Prot (release 2014_11).

### Statistical analyses of metagenomes and metaproteomes

#### MLG-based random forest classification

Relative abundance data of all MLGs were subjected to random forest (RF) analysis to perform five-fold cross validation (R 3.3.2, caret package 6.0-77) [32]. The combinations of optimal MLGs markers maximising the discrimination accuracy between each two groups were thus determined by RF using an embedded feature selection strategy as previously reported [33]. The importance values of model-selected MLGs were calculated using “mean decrease in accuracy” strategy.

#### Spearman’s rank coefficient correlation

Spearman’s rank coefficient correlation (SCC) analysis was used for correlations between MLG profiles and phenotypic factors, and between number of meta-proteins and metagenomic abundances at the genus level, and between the levels of proteins. The significance cut-off for SCC was set at an FDR adjusted *P* < 0.05.

#### Enrichment analysis of KEGG modules

Differentially enriched KEGG modules were identified according to reporter Z-scores [34]. Z-score for each KO was first calculated from Benjamín-Hochberg (BH)-adjusted P values from Wilcoxon rank-sum tests of comparisons between each two groups. The aggregated Z-score for each module was calculated using Z-scores of all individual KOs belonging to the corresponding module. A module was considered significant at a |reporter Z-score | ≥ 1.96.

#### Other statistical analyses

Kruskal–Wallis test was conducted to detect the differences in continuous phenotypic factors, microbial diversity, richness and MLG relative abundances between multi-groups. *Dunn’s post hoc* tests followed by pairwise comparisons were applied to explore the differential phenotypes and MLGs between each two groups (R version 3.3.2, PMCMR package 4.1). The *Dunn’s post hoc* p-values were adjusted with the Benjamini-Hochberg method among multiple pairwise comparisons. The significance cut-off was set as a *Dunn’s post hoc P* value less than 0.05. Wilcoxon rank-sum test was performed for comparisons of MLG relative abundances between published TN-T2D patients from Shanghai [9] and NGT or Pre-DM from the Suzhou cohort in this study for validation purposes. The significance cut-off of Wilcoxon rank-sum test was set as a *P* value less than 0.05. Detailed information on enrichment of MLGs between groups is provided in **Table S5**.

Wilcoxon rank-sum test was conducted to detect differences in protein levels between each two groups. The significance cut-off for proteins was set as a *P* value less than 0.05, and a fold change of protein levels > 1.2 or < 0.8. Chi-square test was conducted to detect the distribution of differences in discrete phenotypic factors, such as sex and treatment distribution between groups, and to identify differences in taxonomic and functional assignments between metagenomic and metaproteomic datasets. The significant cut-off was set as *P* value less than 0.05.

#### Data availability

Metagenomic sequencing data for 254 faecal samples can be accessed from China Nucleotide Sequence Archive (CNSA) with the dataset identifier CNP0000175. The mass spectrometry metaproteomics data have been deposited to the ProteomeXchange Consortium via the PRIDE partner repository with the dataset identifier PXD013452 and 10.6019/PXD013452.

## Results

### Experimental design

The cohort consisted of 77 TN-T2D patients, 80 Pre-DM individuals and 97 NGT individuals from Suzhou, China (**Methods, Table S1**). The three groups were matched regarding body mass index (BMI) and sex (P > 0.05), but individuals with TN-T2D (mean age 66 +/− 8 years) were on average 5 years older than individuals in the two other groups (**Table S1**). Shotgun metagenomics was performed on faecal samples from all participants, whereas metaproteomics profiling was performed on a subgroup of 84 participants, including 28 age-, sex-, and BMI-matched individuals from each group (**Figure 1**).

**Figure 1.**
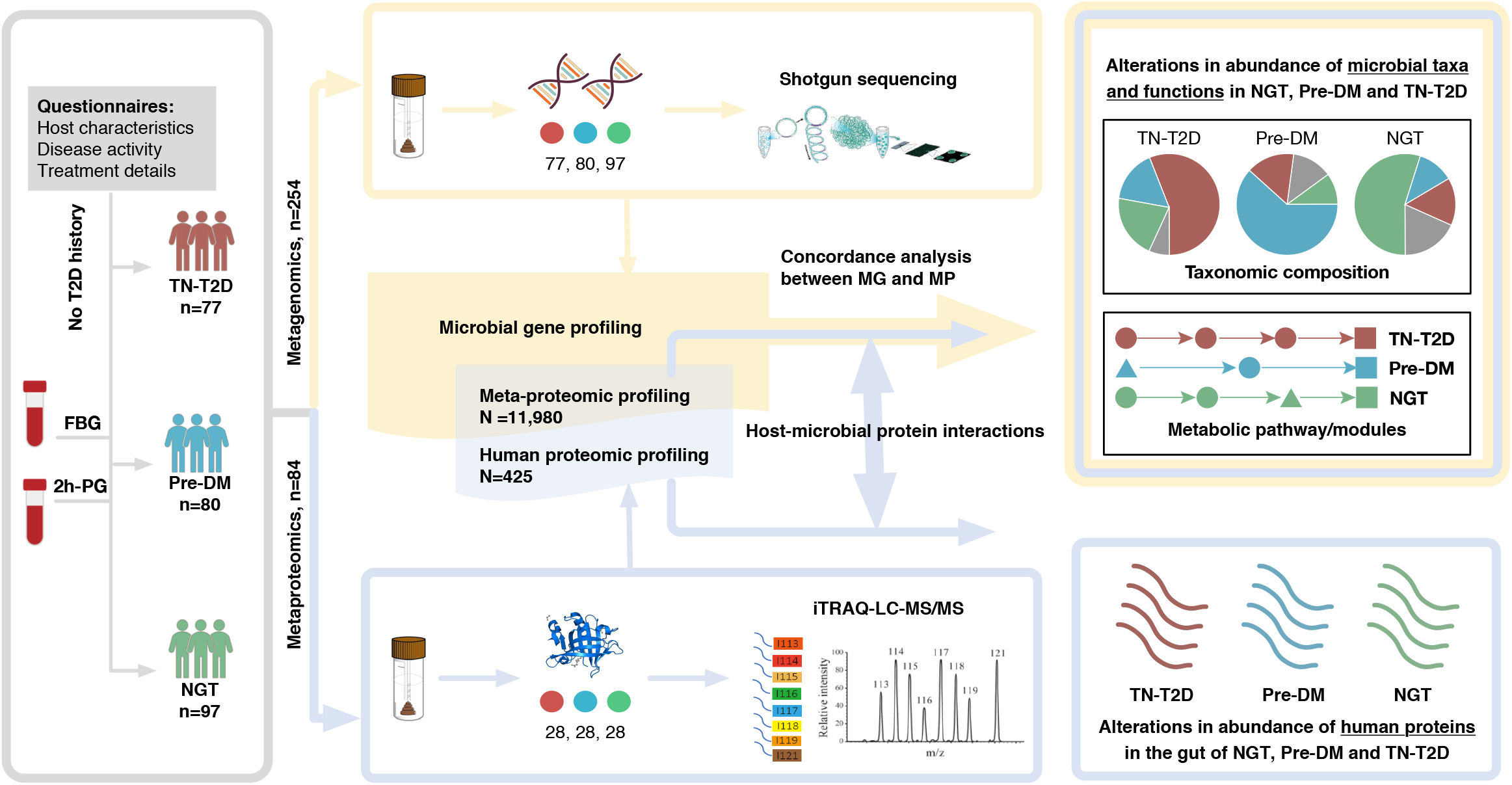
Experimental overview. 254 participants were recruited from the Suzhou cohort and diagnosed as treatment naive T2D patients (TN-T2D, n=77, red), prediabetic individuals (Pre-DM, n=80, blue) or individuals with normal glucose tolerance (NGT, n=97, green). Each participant provided two stool samples. One set of stool samples was used for metagenomic shotgun sequencing, followed by IGC-based taxonomic and functional analyses. The other set of stool samples, comprising a total of 84 samples with 28 age-, BMI- and sex-matched participants from each group, was selected for metaproteomic analyses using isobaric tags for relative and absolute quantitation (iTRAQ)–coupled-liquid chromatography tandem mass spectrometry (iTRAQ-LC-MS/MS) to provide information on the microbial and host proteins present in stool samples. A total of 11, 980 meta-proteins and 425 human proteins were identified in this study. Microbial gene and protein profiling were used to determine alterations in the abundance of microbial taxa and functions, and human protein profiling was used to identify alterations in the abundance of human proteins in faecal samples from NGT, Pre-DM and TN-T2D individuals.

### Distinct metagenomics profiles in Chinese prediabetic and type 2 diabetic individuals

Shotgun metagenomic sequencing of the 254 stool DNA samples was performed using the BGISEQ-500 platform and raw reads were filtered and aligned to the integrated gene catalogue (IGC) of the human gut microbiome to generate gene, taxonomic and functional profiles as previously described (**Methods, Table S2**). In line with previous studies [4–6], no significant differences in microbial gene-based richness, alpha-diversity, and beta-diversity were found between the NGT, Pre-DM, and TN-T2D individuals (**Figure S3**, Kruskal-Wallis (KW) test, *P* > 0.05). Using a metagenome-wide association approach [4], we identified 266,015 T2D-associated genes (KW test, *P* < 0.05) and clustered these genes into 126 metagenomic linkage groups (MLGs, ≥100 genes, **Table S5**).

We further applied the KW test to detect statistically significant differences in the relative abundances of MLGs between individuals with NGT, Pre-DM, and TN-T2D. Compared to NGT individuals, the abundances of MLGs from the *Clostridia* class, such as *Butyrivibrio crossotus* (MLG-2076), *Dialister invisus* (MLG-3376) and *Roseburia hominis* (MLG-14865 and MLG-14920) were significantly lower in individuals with Pre-DM or TN-T2D (**Figure 2A, Table S5**, *Dunn’s post hoc test, P* < 0.05), which is in agreement with previous findings in a Danish T2D cohort [6]. In addition, we found that the abundance of the butyrate-producing *Faecalibacterium prausnitzii* (MLG-4560) was lower in Pre-DM compared to both NGT and TN-T2D individuals. On the contrary, MLGs annotated to *Escherichia coli* (MLG-7919 and MLG-7840), *Streptococcus salivarius* (MLG-6991 and MLG-7099), and *Eggerthella sp*. (MLG-351) were highly enriched in Pre-DM compared to NGT individuals (**Figure 2A**, *P* < 0.05). An increased abundance of *Streptococcus* operational taxonomic units (OTUs) was also recently reported in a Danish prediabetic cohort [7]. Additionally, Pre-DM individuals also exhibited a significant enrichment in *E. coli* abundance compared to TN-T2D individuals (**Figure 2A**, *P* < 0.05). Moreover, we detected significantly lower abundances of *Akkermansia muciniphila* (MLG-2159) and *Clostridium bartlettii* (MLG-7540) and higher abundances of *Bacteroides caccae* (MLG-10234 and MLG-10325), *Bacteroides finegoldii* (MLG-10154 and MLG-10159), and *Collinsella intestinalis* (MLG-10084) in TN-T2D patients compared with NGT and Pre-DM individuals (**Figure 2A**, *P* < 0.05). Finally, the abundance of *Megasphaera elsdenii* (MLG-1568) was significantly higher in both TN-T2D and Pre-DM individuals than in NGT individuals (**Figure 2A**, *P* < 0.05), in line with the positive correlation between the relative abundance of the genus *Megasphaera* and T2D recently reported in a large cohort with about 7000 individuals from South China [35]. Several key findings were further validated in faecal samples of 94 treatment naïve T2D patients in Shanghai (Gu et al., 2017a), such as a lower abundance of *A. muciniphila* and *C. bartlettii* compared to NGT and Pre-DM individuals, and a lower abundance of *E.coli* compared to Pre-DM individuals in this study (**Figure 2A, Table S5**, *Wilcoxon* rank test, *P* < 0.05). A summary of gut microbial taxa reported in previously published cross-sectional T2D or prediabetes studies is presented in **Table S6**.

**Figure 2.**
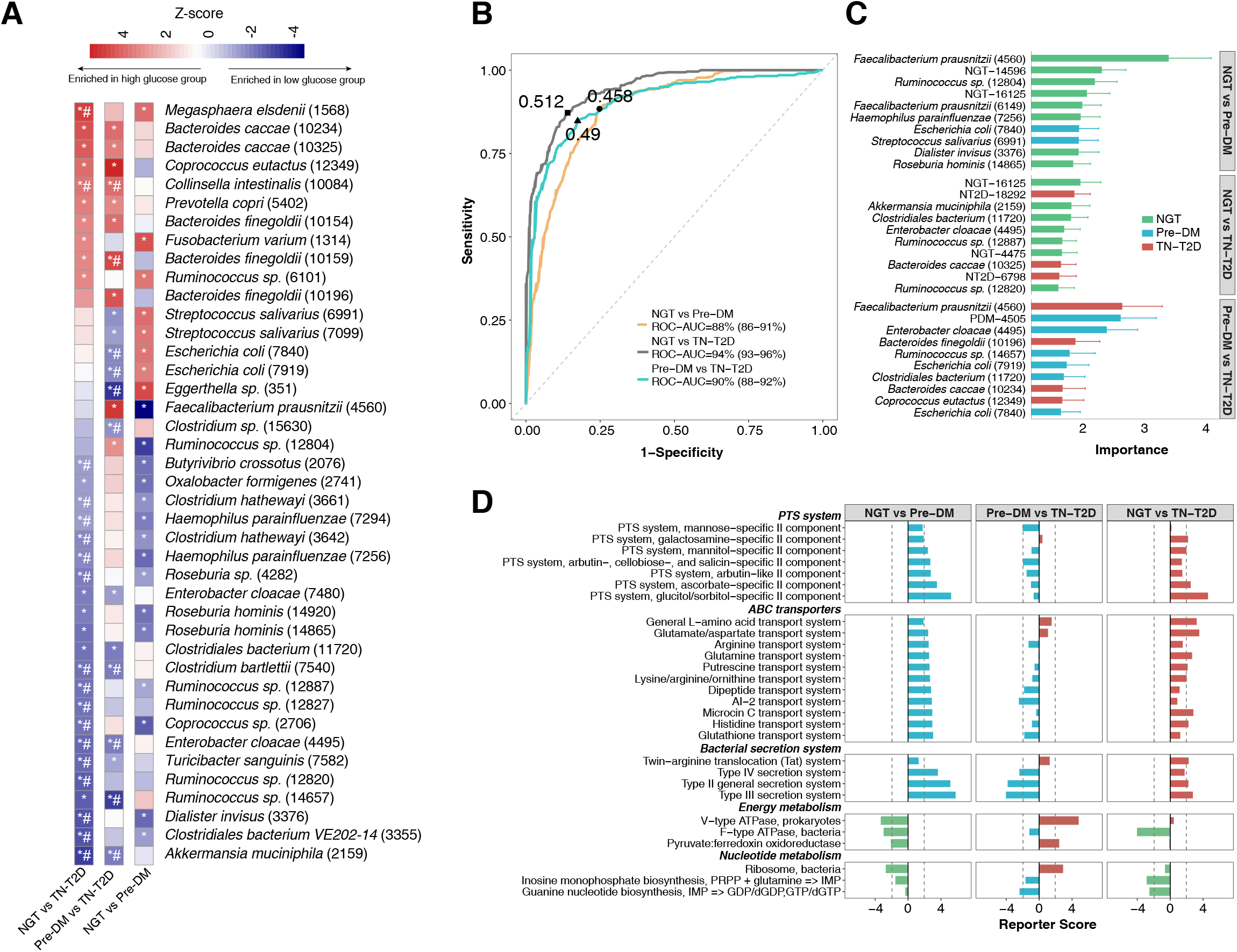
Determination of alterations in the abundance of MLGs and functional modules. **(A)** Heatmap of statistically significant annotated MLGs discriminating between TN-T2D, Pre-DM and NGT based on Z-scores. Red, MLGs enriched in high glucose groups, blue, MLGs enriched in low glucose groups. *, indicates MLGs significantly differed between any two groups in the Suzhou cohort; *Dunn’s post hoc* test, *P* < 0.05. #, indicates significant MLGs replicated in the treatment naïve T2D patients from Shanghai (Gu et al., 2017a) compared with Pre-DM and NGT in the Suzhou cohort; Wilcoxon rank-sum test, *P* < 0.05 (See **Table S5** for full list). **(B)** Performance of cross-validated random forest (RF) classification models using relative abundance profiles of gut microbial MLGs, assessed by the area under the ROC curve (AUC), 95% confidence intervals (CI). Orange, AUC for the RF model classifying NGT (n=97) and Pre-DM (n=80). Grey, AUC for the RF model classifying NGT (n=97) and TN-T2D (n=77). Blue, AUC for the RF model classifying Pre-DM (n=80) and TN-T2D (n=77). The best cut-off points are marked on the ROC curves. **(C)** Bar plot showing the 10 most discriminating MLGs in the RF models for distinguishing between NGT, Pre-DM and TN-T2D. The bar lengths indicate the importance of the selected MLGs, and colours represent enrichment in NGT (green), Pre-DM (blue) and TN-T2D (red). **(D)** Differential enrichment of KEGG modules comparing TN-T2D, Pre-DM and NGT. Dashed lines indicate a reporter score of 1.96, corresponding to 95% confidence in a normal distribution.

We next performed Spearman’s rank correlation analysis to explore the associations between host phenotypes and MLGs. *M. elsdenii* and four unannotated MLGs enriched in TN-T2D individuals showed significantly positive correlations to glycaemic indices, including homeostasis model assessment of insulin resistance (HOMA-IR), fasting blood glucose (FBG), 2h post-load glucose (2h-PG), and HbA1c, whereas MLGs enriched in NGT were negatively correlated with the abovementioned indices (adjusted *P* < 0.05, **Figure S4A-B**). Very few MLGs showed significant correlations with non-glycaemic indices, such as age, BMI and systolic blood pressure (SBP) (**Figure S4**).

To assess the discriminative power of MLGs in T2D and identify key MLGs differentiating individuals with respect to different disease stages, we applied a feature selection approach and constructed Random Forest (RF) classification models comparing the groups (**Methods**). Remarkably, the RF models provided high performances regarding classification of samples from the two different disease stages, with area under the ROC curve (AUC) values from 0.90 to 0.94 (**Figure 2B**). Apart from taxonomically unclassified MLGs, the most discriminatory MLG for separating TN-T2D and NGT was *A. muciniphila*. Moreover, MLGs annotated to *F. prausnitzii* and *E. coli* both showed to be important in separating Pre-DM samples from TN-T2D and NGT samples (**Figure 2C**), indicating the unique microbial signatures of lower abundance of *F. prausnitzii* and higher abundance of *E. coli* in Pre-DM individuals. We also validated the predictive power of the RF models between TN-T2D and other two groups, which showed an accuracy of 76. 6% (72 of 94 patients) for disease prediction in a previously described TN-T2D cohort from Shanghai (**Table S7**) [9].

We next performed KEGG enrichment analyses to examine possible differential patterns of microbial functional potentials in NGT, Pre-DM and TN-T2D individuals (**Table S8**). Interestingly, we observed a significant enrichment in modules comprising several sugar phosphotransferase systems (PTS), ATP-binding cassette transporters (ABC transporters) of amino acids, and bacterial secretion systems in the gut microbiota of Pre-DM compared to NGT individuals (reporter score ≥ 1.96, **Figure 2D**). Likewise, in line with previous findings in several Chinese cohorts with metabolic diseases, such as atherosclerotic cardiovascular disease (ACVD), obesity and T2D [36], a similar enrichment was found in TN-T2D patients compared with NGT individuals (**Figure 2D**). The abundances of the transport system for microcin C, a peptide-nucleotide antibiotic produced by *Enterobacteria* [37], and the transport system for autoinducer-2 (AI-2), a quorum sensing signalling molecule reported in Proteobacteria [38], were also significant higher in Pre-DM than in NGT individuals (**Figure 2D**). Except for enrichment of type II-IV secretion and AI-2 transport systems in Pre-DM vs TN-T2D, we found no other KEGG modules for PTS and ABC transporters to differ significantly in abundance between Pre-DM and TN-T2D individuals (**Figure 2D**). However, Pre-DM individuals displayed a significant reduction with respect to several energy and nucleotide metabolism modules compared to both NGT and TN-T2D individuals, including modules of V-type ATPase, pyruvate: ferredoxin oxidoreductase, and bacterial ribosomal proteins (**Figure 2D**). Taken together, these results indicate the possible involvement of substantial compositional and functional disease-related gut microbial changes in the pre-diabetic stage.

### Gut metaproteomics simultaneously identifies faecal levels of microbial and human proteins

To gain further insights into functional changes in the gut microbiota associated with T2D, we conducted metaproteomic analyses using iTRAQ (isobaric peptide tags for relative and absolute quantification) and LC-MS/MS-based protocols on 84 samples, with 28 samples derived from each of the three diagnostic groups (**Methods, Figure S1**). Using the strict parameters of 2 peptide-spectrum matches (PSMs) per protein, < 10 ppm mass error and 1% PSM-level FDR (**Methods**), we identified a total of 145,014 high quality PSMs corresponding to 15,670 proteins, including 15,245 (97.29%) microbial proteins and 425 (2.71%) human proteins (**Table S9**). As reported [14,19,20], one microbial peptide often exhibits matches to multiple proteins with high sequence similarity, resulting in difficulties in identifying the microbial origin of individual peptides. To alleviate ambiguities, we applied a maximum parsimony principle reported in recent studies [14], [39] and generated 11,980 non-redundant meta-proteins (78.58% of microbial proteins) containing at least one unique microbial peptide. The relative intensities of these unique peptides were further used for meta-protein quantification (**Methods, Table S9**). The number of identified meta-proteins ranged between 5,067 in the Pre-DM samples to 8,134 in the TN-T2D samples (**Table S9**). Venn diagrams showed that only 2782 meta-proteins (34.2%-54.9% of the total number of meta-proteins per group) were shared among the three groups (**Figure S5A**), indicating differential microbial expression patterns at the protein level among the groups. Taxonomic annotations indicated a higher percentage of unique Proteobacteria meta-proteins in Pre-DM individuals, compared to the other groups (Chi-square test, *P* < 0.05, **Figure S5B**), whereas no difference in the distributions of the uniquely detected meta-proteins associated with a wide range of functions was found between the three groups (**Figure S5C**).

### Concordance and discordance of microbiota features between metagenomes and metaproteomes

Based on annotated microbial features, we next investigated the consistency as well as the divergence of microbial composition and function at the DNA and protein level. At the phylum level, more than 90% genes and meta-proteins were consistently assigned to three major phyla, namely Firmicutes, Bacteroidetes and Proteobacteria (**Figure 3A**). Despite the overall consistency, we found a significantly higher percentage of the annotated proteins to be assigned to Bacteroidetes (41%) compared to the percentage of genes annotated to Bacteroidetes (25%) (Chi-square test, *P* < 0.05, **Figure 3A**), suggesting that Bacteroidetes might display an overall higher protein production than the other phyla across the 84 samples. At the genus level, the composition of the metaproteomes was biased towards a limited number of genera. Among 212 common metagenomically-identified genera detected in at least 10% of the 84 samples, only 81 genera (38.21%) could be detected based on metaproteomics (**Table S10**). Spearman’s rank correlation analysis was subsequently performed to determine the relationship between the number of meta-proteins and the abundances at the genus level based on metagenomics. The more abundant a given genus was based on metagenomics analysis, the more of the identified meta-proteins were assigned to this genus (Spearman’s correlation coefficient (SCC) = 0.726, *P* = 5.21E-08, **Figure 3B, Table S9**), with *Bacteroides* (n=1664), *Prevotella* (n=818) and *Faecalibacterium* (n=719) harbouring most assigned meta-proteins. For a few genera, such as *Anaerotruncus* (n=9), *Paraprevotella* (n=9) and *Enterococcus* (n=7), we were only able to identify less than 10 meta-proteins although their median metagenomic abundances were greater than 1E-04 (**Table S10**).

**Figure 3.**
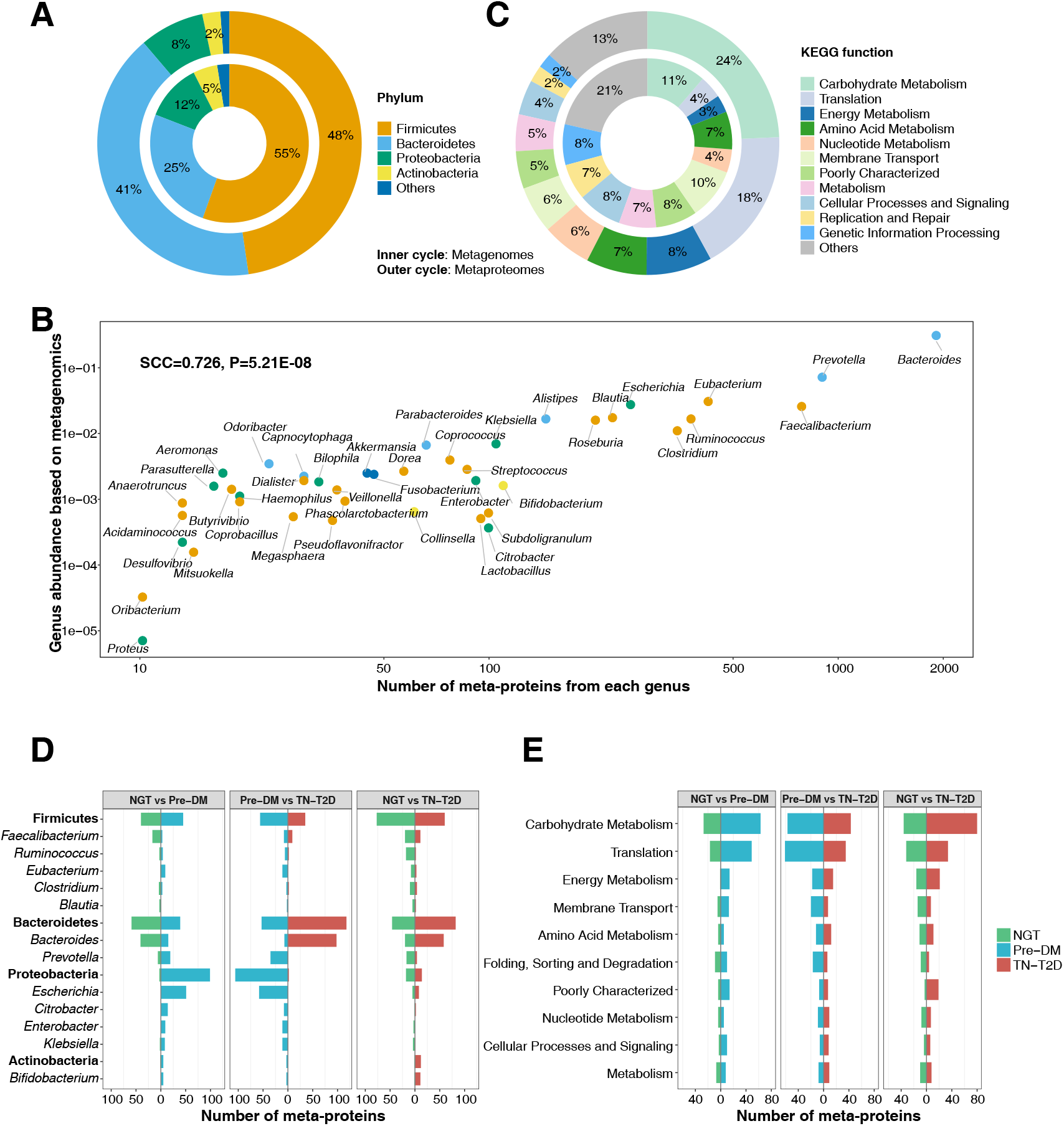
Concordance and discordance of gut microbiome features in metagenomes and metaproteomes. **(A)** Taxonomic distribution at the phylum level. Inner circle, metagenomes; Outer circle, metaproteomes. **(B)** Spearman’s rank correlation between the median relative abundances of genera in metagenomes of 84 samples selected for metaproteomics and the number of identified meta-proteins assigned to the same genus. (C) Functional distribution at KEGG level 2. Inner circle, metagenomes; Outer circle, metaproteomes. **(D-E)** Enrichment analysis of differentially expressed meta-proteins at taxonomic (d) and functional levels (e) comparing NGT, Pre-DM and TN-T2D individuals. The number of meta-proteins that exhibited significant differences in levels in each pairwise comparison is shown. Colours represent enrichment in NGT (green), Pre-DM (blue) and TN-T2D (red). Significant enrichment is defined as *P* < 0.05 (Wilcoxon rank-sum test) with a fold change of mean intensities > 1.2 in pairwise comparisons.

Comparing KEGG functional categories based on metagenomics and metaproteomics data, we observed large differences in the relative contribution of individual categories between the two datasets (Chi-square test, *P* < 0.05, **Figure 3C**), in accordance with several previous studies [14,19,20]. For instance, as determined by metaproteomics, 24% and 18% of the proteins were assigned to carbohydrate metabolism and translation categories, whereas the corresponding metagenomic percentages of the two categories were only 11% and 4%, respectively (**Figure 3C**). We found that 1508 meta-proteins, accounting for 12.59% of all identified meta-proteins, could be assigned to 10 KEGG orthologues (KO). The top KOs harboured 360 proteins annotated as Ca-activated chloride channel homologues (K07114), whereas the remaining KOs comprised proteins representing abundant house-keeping proteins such as elongation factors, large subunit ribosomal proteins (K02355, K02358 and K02395), chaperones (K04077 and K04043), and glyceraldehyde 3-phosphate dehydrogenase (K00134) as well as flagellin proteins (K02406) (**Table S11, Figure S6A**).

Aiming to link the microbial protein patterns to metagenomic microbial abundances, we next conducted a fold-change analysis of meta-proteins. In agreement with our metagenomic findings (**Figure 2A**), the Proteobacteria meta-proteins (mainly from *Escherichia, Citrobacter* and *Enterobacter)* exhibited enrichment in the Pre-DM group, whereas *Bacteroides* meta-proteins were enriched in TN-T2D individuals (**Figure 3D, Table S12**, *P* < 0.05 and fold change of protein intensities > 1.2). Surprisingly, *Prevotella* meta-proteins were selectively enriched in Pre-DM individuals (**Figure 3D**), although no *Prevotella* annotated metagenomic MLGs exhibited significantly higher abundance. At the functional level, we observed that the level of meta-proteins involved in carbohydrate metabolism tended to be lower in NGT compared to Pre-DM and TN-T2D individuals, including those involved in the metabolism of succinate (**Figure 3E, Figure S6B, Table S11**).

### Functional characteristics of faecal excreted human proteins in T2D

Among the 425 detected human proteins, we identified 218 human proteins that were shared among the NGT, Pre-DM, and TN-T2D groups, accounting for 59.6% to 85.2% of the identified human proteins in each group (**Figure S7A**). We next annotated the human proteins with Gene Ontology (GO) terms to obtain insight into the functional characteristics of the human proteins excreted in faeces (**Table S13**). Among the identified proteins, 181 (42.59%) had previously been identified in faecal samples by metaproteomics, indicative of their general presence (**Table S14**) [14,19,20]. These included several intestinal mucin proteins, such as MUC-1, MUC-2, MUC-4, MUC5B, MUC12 and MUC-13 as well as members of annexins (ANXA1-ANXA7, a family of calcium-binding proteins) (**Table S14**). We identified 233 of the faecal human proteins to have tissue-specific annotation, amongst which 151 proteins (64.81%) were reported to exhibit high expression in the digestive system, and the remaining proteins were annotated to be highly expressed in blood or other tissues such as epidermis (**Table S13**). Of interest, 18 of the human proteins were annotated as AMPs [40] (**Table S13**). Several human proteins involved in glucose metabolism, including the sodium/glucose cotransporter 1, were detected in faecal samples of TN-T2D patients only (**Figure S6B**). Inhibitors of this protein have been proposed for antidiabetic treatment ^26^. Additionally, the TMAO-producing enzyme, dimethylaniline monooxygenase [N-oxide-forming] 3 (FMO3) was also identified exclusively in the TN-T2D group (**Table S13**). On the other hand, we found that ras GTPase-activating-like protein (IQGAP1) and unconventional myosin-Ic (MYO1C) were uniquely identified in the NGT group (**Figure S7B**). Loss of IQGAP1 and MYO1C has been related to impairment of insulin signalling [43–45], but whether their presence in faeces has functional implications remains to be established.

Forty-nine of the human proteins present in faeces were found to differ significantly in intensity between at least two of the groups (**Figure 4A, Table S15**). We found significantly higher levels of four AMPs, including defensin-5, neutrophil defensin-1, lysozyme c, as well as secreted phospholipase A2, all with important roles in the defence against bacteria [46–48], in faecal samples from NGT individuals than in samples from TN-T2D individuals (**Figure 4A**). We also found higher levels of mucin-5AC samples from NGT compared to TN-T2D individuals, suggesting possible effects on the mucus barrier in TN-T2D. Interestingly, the level of the antimicrobial cathepsin G, reported to inhibit the growth of several organisms from the Proteobacteria phylum [49], was higher in samples from Pre-DM than NGT and TN-T2D, and this was coupled to lower levels of alpha-1-antichymotrypsin and alpha-1-antitrypsin, both known inhibitors of cathepsin G [50] (**Figure 4A**), suggesting that Pre-DM individuals have initiated strategies to activate a defence system against the enhanced relative abundances of E. *coli*. By contrast, we found that several proteins within the immunoglobulin superfamily were present at lower levels in samples from Pre-DM compared to NGT or TN-T2D (**Figure 4A**). Individuals with Pre-DM also exhibited lower levels of galectin-3, a lectin with beta-galactoside-binding ability. Galectin-3 has been reported to bind lipopolysaccharides (LPS) from *E. coli* and play a role as a negative regulator of LPS-mediated inflammation [51]. In addition, galectin-3 was also reported to improve epithelial intercellular contact via desmoglein-2 stabilization [52]. Taken together, these finding indicate that the gut ecosystem in Pre-DM individuals exhibits trait compatible with the upregulation of defence systems against an increased abundance of Proteobacteria simultaneously with the downregulation of factors capable of reducing the impact of the inflammation-inducing activity of LPS. We also found that several digestive enzymes differed in levels in faeces from NGT, Pre-DM, and TN-T2D individuals. Thus, we found lower levels of proteases (trypsin and chymotrypsin and their precursors) and lipases, and higher amylase (AMY1) levels in TN-T2D (Figure 4A). It is also interesting to note that the level of dipeptidyl peptidase 4 (DDP4), known to inhibit insulin secretion via its action on GLP-1, was lower in individuals with Pre-DM than in TN-T2D individuals. A network analysis revealed significant correlations between 20 human proteins showing significant differences in levels in two-pairwise comparisons between NGT, Pre-DM and TN-T2D individuals (**Figure 4B**). For instance, we identified a negative correlation between the defensin-5 and TN-T2D-enriched peptidyl-prolyl cis-trans isomerase B (PPIB) (**Figure 4B**, SCC, adjusted *P* < 0.05), the latter previously reported to be associated with islet dysfunction [53].

**Figure 4.**
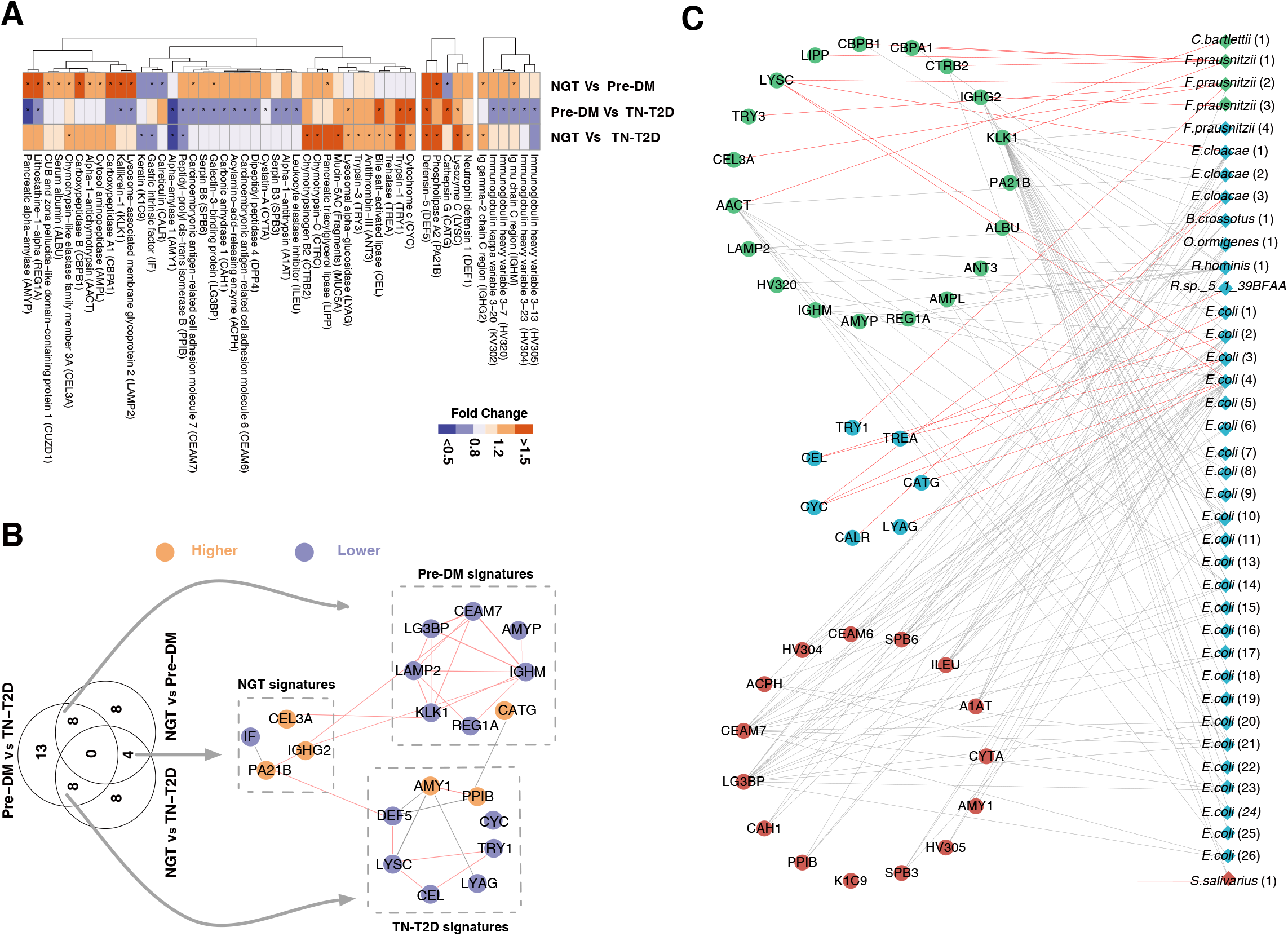
Characterisation of human proteins in faecal samples from Chinese NGT, Pre-DM, and TN-T2D individuals. **(A)** Heatmap showing levels of 49 discriminatory human proteins as fold change between each two groups. *, *P* < 0.05 and fold change of protein levels > 1.2 or < 0.8. **(B)** Protein-protein interaction network based on 20 discriminatory human proteins in at least two pair-wise comparisons. The group signatures indicate human proteins with significantly higher or lower levels in this group compared to others. Orange indicates higher protein levels and blue indicates lower protein levels. **(C)** Protein-protein interactions based on discriminatory meta-proteins in pair-wise comparisons and discriminatory human proteins. Only discriminatory meta-proteins annotated to the corresponding taxon of the MLGs were selected for the analysis. The circles indicate human proteins and diamonds indicate meta-proteins. Detailed information on the numbered meta-proteins is presented in **Table S12**. Colours represent protein enrichment in NGT (green), Pre-DM (blue) and TN-T2D (red). Pink line indicates positive correlation and grey line indicates negative correlation (Spearman’s rank correlations, adjusted *P* < 0.05).

Aiming to investigate possible host-microbial protein interactions in the human gut, we next investigate the possible correlation between the discriminatory bacterial and human proteins. Interestingly, we found significantly negative correlations between several Pre-DM-enriched *E. coli* proteins and human proteins involved in innate immune responses (HV304, HV305) and adhesion (CEAM6, CEAM7), whereas positive correlations were found between *E. coli* proteins and cathepsin G, Cytochrome c (CYC) and trypsin–1 (TRY1) (**Figure 4C**, adjusted *P* < 0.05). Conversely, NGT-enriched proteins from *F. prausnitzii* showed positive correlations with several NGT-enriched digestive enzymes from the exocrine pancreas, such as chymotrypsin-like elastase family member 3A (CEL3A), chymotrypsinogen B2 (CTRB2) and carboxypeptidases (CBPA1 and CBPB1).

## Discussion

Our comparative study using metagenomics and metaproteomics in normal glucose tolerant, pre-diabetics and treatment naïve T2D individuals provides important novel findings with regard to disease-stage specifications at the gut bacterial and host level. A substantial number of Pre-DM associated features were revealed at both the metagenomics and metaproteomics level. Of specific note are the significantly higher abundance of Proteobacteria species (dominated by *E. coli*) and the lower levels of host proteins which potentially are involved in Proteobacteria-specific responses in Pre-DM, such as galectin-3 and proteins within the immunoglobulin superfamily. Furthermore, significantly higher levels of *Prevotella* proteins were uniquely detected in Pre-DM individuals although the abundance of *Prevotella* was not significantly enriched in this group based on metagenomics data. *Prevotella copri* has previously been shown to produce branched-chain amino acids (BCAA), reported to correlate with BCAA blood levels and insulin resistance [54]. However, in the present study only two enzymes related to the synthesis of BCAAs were detected among the identified *Prevotella* proteins with no differences in levels between the three groups.

Only a modest number of relatively highly abundant faecal proteins were identified in the current study. This reflects the current methodological challenges in microbial protein extraction, identification, and annotation as reported previously [55,56], as well as the detection limitations of MS-based proteomics [57]. For instance, we identified less than 50 proteins from each of several taxa with median abundances in the 0.1 % ranges based on metagenomics data (such as NGT-enriched *Dialister*, *Butyrivibrio* and *Haemophilus*). Nevertheless, metaproteomics provides a valuable addition to not only estimating expression of microbial proteins, but also to delineate host-microbial protein interactions in different disease stages. In this regard, we identified higher levels of several host-derived AMPs in NGT individuals compared to TN-T2D and Pre-DM individuals, suggesting a possible stronger host defence against invading (disease-related) microbes in NGT individuals. By contrast, significant negative associations were found between Pre-DM-enriched *E. coli* proteins and several human proteins, including AMPs, adhesion molecules and galectin-3, all involved in intestinal barrier function. It is also worth to note the significant changes in levels and types of digestive enzymes identified in the faecal samples, where TN-T2D showed enhanced alpha-amylase (AMY1) levels, as compared to pancreatic-derived lipases and proteases. However, the level of pancreatic alpha-amylase (AMYP) was lower in Pre-DM compared to the two other groups. A metaproteomics study has reported lower faecal AMYP levels in type 1 diabetes (T1D) patients compared to their healthy relatives ^10^, whereas no difference in levels of AMY1 was reported between T1D and controls, suggesting different amylase responses might be present in Pre-DM, TN-T2D and T1D patients based on metaproteomics data. Differences in levels of secreted digestive enzymes from the exocrine pancreas in NGT, Pre-DM and T2D have to our notice not been addressed previously, although it may be of major importance in relation to the metabolic state in T2D.

Together, our findings suggest that unique and nonlinear changes of the intestinal ecosystem might exist in Pre-DM individuals before transition to T2D. Further large-scale, longitudinal follow-up studies are needed to delineate how microbial functions changes from prediabetes to diabetes and to address the nature of interactions between the gut microbiota and the host in the transitional phases leading to overt T2D.

## Acknowledgements

We thank Prof. Yan Ren for helpful discussion on designing the metaproteomic experiments. We thank Dr Cong Lin and Dr Zhe Zhang for helpful discussion and suggestions on developing the manuscript. We gratefully acknowledge colleagues at BGI for DNA extraction, library preparation and shotgun sequencing experiments, and helpful discussions.

## Author contributions

J.L. and H.Z. designed and coordinated the study. F.L. and J.Z. oversaw the blood and faecal sample collection. Y.L, B.C., J.C., X. B., Y.H. and Y.G. participated in sample collection and provided phenotypic information. G.H., B.Z, J.Z. and S.L. carried out the metaproteomic experiments. H.Z., H.R., F.Y., Z.S, and H.Zou. performed the bioinformatic analyses of metagenomic data. H.Z., H.R., C.F., B.Z, G.H., Y.Z. and J.W. performed the bioinformatic analyses of metaproteomic data. H.Z. and H.R performed integrative analyses of metagenomic and metaproteomic data. Y.Z. performed revision of the figures. H.Z. interpreted together with J.L., S.B. and K.K. the data and wrote the first version of the manuscript. J.L., K.K., S.B., and L.M. performed revision of the manuscript. H.Z., H.R., C.F., G.H., F.Y., Z.Y., Y.Z., Z.S., J.W, L.M., S.B., K.K. and J.L. participated in discussions. All authors contributed to the revision of the manuscript. All authors read and approved the final manuscript.

## Reference

[1] Stumvoll M, Goldstein BJ, Van Haeften TW. Type 2 diabetes: Principles of pathogenesis and therapy. Lancet, vol. 365, 2005, p. 1333–46. doi:10.1016/S0140-6736(05)61032-X.

[2] Pickup JC. Inflammation and Activated Innate Immunity in the Pathogenesis of Type 2 Diabletes. Diabetes Care 2004;27:813–23. doi:10.2337/diacare.27.3.813.

[3] Wang L, Gao P, Zhang M, Huang Z, Zhang D, Deng Q, et al. Prevalence and ethnic pattern of diabetes and prediabetes in China in 2013. JAMA - J Am Med Assoc 2017;317:2515–23. doi:10.1001/jama.2017.7596.

[4] Wang J, Qin J, Li Y, Cai Z, Li S, Zhu J, et al. A metagenome-wide association study of gut microbiota in type 2 diabetes. Nature 2012;490:55–60. doi:10.1038/nature11450.

[5] Karlsson FH, Tremaroli V, Nookaew I, Bergström G, Behre CJ, Fagerberg B, et al. Gut metagenome in European women with normal, impaired and diabetic glucose control. Nature 2013. doi:10.1038/nature12198.

[6] Forslund K, Hildebrand F, Nielsen T, Falony G, Le Chatelier E, Sunagawa S, et al. Disentangling type 2 diabetes and metformin treatment signatures in the human gut microbiota. Nature 2015;528:262–6. doi:10.1038/nature15766.

[7] Allin KH, Tremaroli V, Caesar R, Jensen BAH, Damgaard MTF, Bahl MI, et al. Aberrant intestinal microbiota in individuals with prediabetes. Diabetologia 2018;61:810–20. doi:10.1007/s00125-018-4550-1.

[8] Wu H, Esteve E, Tremaroli V, Khan MT, Caesar R, Mannerås-Holm L, et al. Metformin alters the gut microbiome of individuals with treatment-naive type 2 diabetes, contributing to the therapeutic effects of the drug. Nat Med 2017;23:850–8. doi:10.1038/nm.4345.

[9] Gu Y, Wang X, Li J, Zhang Y, Zhong H, Liu R, et al. Analyses of gut microbiota and plasma bile acids enable stratification of patients for antidiabetic treatment. Nat Commun 2017;8:1785. doi:10.1038/s41467-017-01682-2.

[10] Zhao L, Chen Y, Xia F, Abudukerimu B, Zhang W, Guo Y, et al. A glucagon-like peptide-1 receptor agonist lowers weight by modulating the structure of gut microbiota. Front Endocrinol (Lausanne) 2018. doi:10.3389/fendo.2018.00233.

[11] Moreira G V., Azevedo FF, Ribeiro LM, Santos A, Guadagnini D, Gama P, et al. Liraglutide modulates gut microbiota and reduces NAFLD in obese mice. J Nutr Biochem 2018. doi:10.1016/j.jnutbio.2018.07.009.

[12] Olivares M, Neyrinck AM, Pötgens SA, Beaumont M, Salazar N, Cani PD, et al. The DPP-4 inhibitor vildagliptin impacts the gut microbiota and prevents disruption of intestinal homeostasis induced by a Western diet in mice. Diabetologia 2018. doi:10.1007/s00125-018-4647-6.

[13] Liao X, Song L, Zeng B, Liu B, Qiu Y, Qu H, et al. Alteration of gut microbiota induced by DPP-4i treatment improves glucose homeostasis. EBioMedicine 2019. doi:10.1016/j.ebiom.2019.03.057.

[14] Heintz-Buschart A, May P, Laczny CC, Lebrun LA, Bellora C, Krishna A, et al. Integrated multi-omics of the human gut microbiome in a case study of familial type 1 diabetes. Nat Microbiol 2016;2. doi:10.1038/nmicrobiol.2016.180.

[15] Abu-Ali GS, Mehta RS, Lloyd-Price J, Mallick H, Branck T, Ivey KL, et al. Metatranscriptome of human faecal microbial communities in a cohort of adult men. Nat Microbiol 2018;3:356–66. doi:10.1038/s41564-017-0084-4.

[16] Liu R, Hong J, Xu X, Feng Q, Zhang D, Gu Y, et al. Gut microbiome and serum metabolome alterations in obesity and after weight-loss intervention. Nat Med 2017;23:859–68. doi:10.1038/nm.4358.

[17] Schirmer M, Franzosa EA, Lloyd-Price J, McIver LJ, Schwager R, Poon TW, et al. Dynamics of metatranscription in the inflammatory bowel disease gut microbiome. Nat Microbiol 2018;3:337–46. doi:10.1038/s41564-017-0089-z.

[18] Bajaj JS, Thacker LR, Fagan A, White MB, Gavis EA, Hylemon PB, et al. Gut microbial RNA and DNA analysis predicts hospitalizations in cirrhosis. JCI Insight 2018;3:1–12. doi:10.1172/jci.insight.98019.

[19] Verberkmoes NC, Russell AL, Shah M, Godzik A, Rosenquist M, Halfvarson J, et al. Shotgun metaproteomics of the human distal gut microbiota. ISME J 2009;3:179–89. doi:10.1038/ismej.2008.108.

[20] Young JC, Pan C, Adams RM, Brooks B, Banfield JF, Morowitz MJ, et al. Metaproteomics reveals functional shifts in microbial and human proteins during a preterm infant gut colonization case. Proteomics 2015;15:3463–73. doi:10.1002/pmic.201400563.

[21] Zhong H, Fang C, Fan Y, Lu Y, Wen B, Ren H, et al. Lipidomic profiling reveals distinct differences in plasma lipid composition in healthy, prediabetic, and type 2 diabetic individuals. Gigascience 2017;6. doi:10.1093/gigascience/gix036.

[22] Fang C, Zhong H, Lin Y, Chen B, Han M, Ren H, et al. Assessment of the cPAS-based BGISEQ-500 platform for metagenomic sequencing. Gigascience 2018;7:1–8. doi:10.1093/gigascience/gix133.

[23] Li J, Jia H, Cai X, Zhong H, Feng Q, Sunagawa S, et al. An integrated catalog of reference genes in the human gut microbiome. Nat Biotechnol 2014;32:834–41. doi:10.1038/nbt.2942.

[24] Zou Y, Xue W, Luo G, Deng Z, Qin P, Guo R, et al. 1,520 reference genomes from cultivated human gut bacteria enable functional microbiome analyses. Nat Biotechnol 2019. doi:10.1038/s41587-018-0008-8.

[25] Austin PC. An introduction to propensity score methods for reducing the effects of confounding in observational studies. Multivariate Behav Res 2011;46:399–424. doi:10.1080/00273171.2011.568786.

[26] Wisniewski JR, Zougman A, Nagaraj N, Mann M. Universal sample preparation method for proteome analysis. Nat Methods 2009;6:359–62. doi:10.1038/nmeth.1322.

[27] Guo J, Ren Y, Hou G, Wen B, Xian F, Chen Z, et al. A Comprehensive Investigation toward the Indicative Proteins of Bladder Cancer in Urine: From Surveying Cell Secretomes to Verifying Urine Proteins. J Proteome Res 2016;15:2164–77. doi:10.1021/acs.jproteome.6b00106.

[28] Pappin DJC, Creasy DM, Cottrell JS. Probability-based Protein Identification by Searching Sequence Databases Using Mass Spectrometry Data. Electrophoresis 1999;20:3551–67.

[29] Elias JE, Gygi SP. Target-Decoy Search Strategy for Mass Spectrometry-Based Proteomics. Proteome Bioinforma 2010:55–71. doi:10.1007/978-1-60761-444-9_5.

[30] Brosch M, Yu L, Hubbard T, Choudhary J. Accurate and sensitive peptide identification with mascot percolator. J Proteome Res 2009;8:3176–81. doi:10.1021/pr800982s.

[31] Wen B, Zhou R, Feng Q, Wang Q, Wang J, Liu S. IQuant: An automated pipeline for quantitative proteomics based upon isobaric tags. Proteomics 2014;14:2280–5. doi:10.1002/pmic.201300361.

[32] Max K, Kuhn M. Building Predictive Models in R Using the caret Package. J Stat Softw 2008. doi:10.1053/j.sodo.2009.03.002.

[33] Tett A, Pasolli E, Farina S, Truong DT, Asnicar F, Zolfo M, et al. Unexplored diversity and strain-level structure of the skin microbiome associated with psoriasis. Npj Biofilms Microbiomes 2017;3. doi:10.1038/s41522-017-0022-5.

[34] Patil KR, Nielsen J. Uncovering transcriptional regulation of metabolism by using metabolic network topology. Proc Natl Acad Sci 2005;102:2685–9. doi:10.1073/pnas.0406811102.

[35] He Y, Wu W, Zheng HM, Li P, McDonald D, Sheng HF, et al. Regional variation limits applications of healthy gut microbiome reference ranges and disease models. Nat Med 2018. doi:10.1038/s41591-018-0164-x.

[36] Jie Z, Xia H, Zhong SL, Feng Q, Li S, Liang S, et al. The gut microbiome in atherosclerotic cardiovascular disease. Nat Commun 2017;8:845. doi:10.1038/s41467-017-00900-1.

[37] Rebuffat S. Microcins in action: amazing defence strategies of Enterobacteria. Biochem Soc Trans 2012;40:1456–62. doi:10.1042/BST20120183.

[38] Pereira CS, Thompson JA, Xavier KB. AI-2-mediated signalling in bacteria. FEMS Microbiol Rev 2013;37:156–81. doi:10.1111/j.1574-6976.2012.00345.x.

[39] Muth T, Behne A, Heyer R, Kohrs F, Benndorf D, Hoffmann M, et al. The MetaProteomeAnalyzer: A powerful open-source software suite for metaproteomics data analysis and interpretation. J Proteome Res 2015;14:1557–65. doi:10.1021/pr501246w.

[40] Wang G, Li X, Wang Z. APD3: The antimicrobial peptide database as a tool for research and education. Nucleic Acids Res 2016;44:D1087–93. doi:10.1093/nar/gkv1278.

[41] Song P, Onishi A, Koepsell H, Vallon V. Sodium glucose cotransporter SGLT1 as a therapeutic target in diabetes mellitus. Expert Opin Ther Targets 2016;20:1109–25. doi:10.1517/14728222.2016.1168808.

[42] Van De Laar FA, Lucassen PL, Akkermans RP, Van De Lisdonk EH, Rutten GE, Van Weel C. α-Glucosidase inhibitors for patients with type 2 diabetes: Results from a Cochrane systematic review and meta-analysis. Diabetes Care 2005;28:154–63. doi:10.2337/diacare.28.1.154.

[43] Rittmeyer EN, Daniel S, Hsu S-C, Osman MA. A dual role for IQGAP1 in regulating exocytosis. J Cell Sci 2008;121:391–403. doi:10.1242/jcs.016881.

[44] Chawla B, Hedman AC, Sayedyahossein S, Erdemir HH, Li Z, Sacks DB. Absence of IQGAP1 protein leads to insulin resistance. J Biol Chem 2017;292:3273–89. doi:10.1074/jbc.M116.752642.

[45] Yip MF, Ramm G, Larance M, Hoehn KL, Wagner MC, Guilhaus M, et al. CaMKII-Mediated Phosphorylation of the Myosin Motor Myo1c Is Required for Insulin-Stimulated GLUT4 Translocation in Adipocytes. Cell Metab 2008;8:384–98. doi:10.1016/j.cmet.2008.09.011.

[46] Wiesner J, Vilcinskas A. Antimicrobial peptides: The ancient arm of the human immune system. Virulence 2010;1:440–64. doi:10.4161/viru.1.5.12983.

[47] Vidarsson G, Dekkers G, Rispens T. IgG subclasses and allotypes: From structure to effector functions. Front Immunol 2014;5. doi:10.3389/fimmu.2014.00520.

[48] Nevalainen TJ, Graham GG, Scott KF. Antibacterial actions of secreted phospholipases A2. Review. Biochim Biophys Acta - Mol Cell Biol Lipids 2008;1781:1–9. doi:10.1016/j.bbalip.2007.12.001.

[49] MacIvor DM, Shapiro SD, Pham CT, Belaaouaj A, Abraham SN, Ley TJ. Normal neutrophil function in cathepsin G-deficient mice. Blood 1999;94:4282–93.

[50] Duranton J, Adam C, Bieth JG. Kinetic mechanism of the inhibition of cathepsin G by α1-antichymotrypsin and α1-proteinase inhibitor. Biochemistry 1998;37:11239–45. doi:10.1021/bi980223q.

[51] Li Y, Komai-Koma M, Gilchrist DS, Hsu DK, Liu F-T, Springall T, et al. Galectin-3 Is a Negative Regulator of Lipopolysaccharide-Mediated Inflammation. J Immunol 2008;181:2781–9. doi:10.4049/jimmunol.181.4.2781.

[52] Jiang K, Rankin CR, Nava P, Sumagin R, Kamekura R, Stowell SR, et al. Galectin-3 regulates desmoglein-2 and intestinal epithelial intercellular adhesion. J Biol Chem 2014;289:10510–7. doi:10.1074/jbc.M113.538538.

[53] Lu H, Yang Y, Allister EM, Wijesekara N, Wheeler MB. The Identification of Potential Factors Associated with the Development of Type 2 Diabetes. Mol Cell Proteomics 2008;7:1434–51. doi:10.1074/mcp.M700478-MCP200.

[54] Pedersen HK, Gudmundsdottir V, Nielsen HB, Hyotylainen T, Nielsen T, Jensen BAH, et al. Human gut microbes impact host serum metabolome and insulin sensitivity. Nature 2016;535:376–81. doi:10.1038/nature18646.

[55] Wilmes P, Heintz-Buschart A, Bond PL. A decade of metaproteomics: Where we stand and what the future holds. Proteomics 2015;15:3409–17. doi:10.1002/pmic.201500183.

[56] Heyer R, Schallert K, Zoun R, Becher B, Saake G, Benndorf D. Challenges and perspectives of metaproteomic data analysis. J Biotechnol 2017;261:24–36. doi:10.1016/j.jbiotec.2017.06.1201.

[57] Schubert OT, Röst HL, Collins BC, Rosenberger G, Aebersold R. Quantitative proteomics: Challenges and opportunities in basic and applied research. Nat Protoc 2017;12:1289–94. doi:10.1038/nprot.2017.040.

