## Supplementary information of FigureS1-S7 for "Distinct gut metagenomics and metaproteomics signatures in prediabetics and treatment-naïve type 2 diabetics"

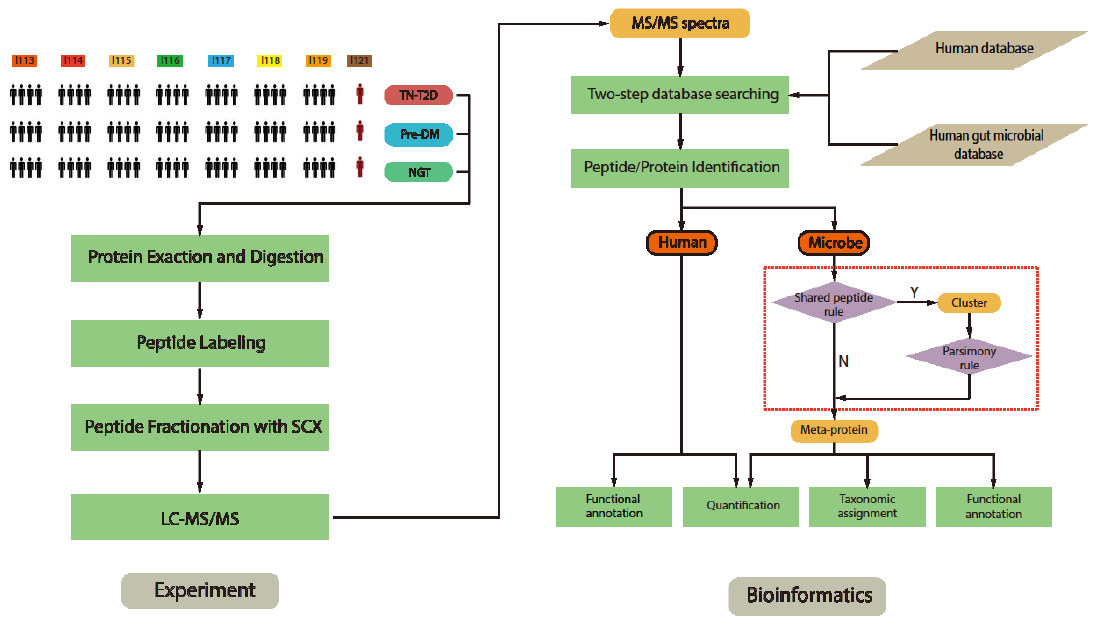

**Figure S1: Flowchart of experimental design and bioinformatic analysis for metaproteomics**

Experimental design: 84 Faecal samples from 28 age-, sex- and BMI-matched individuals from each of the three diagnostic groups were randomly selected for metaproteomic analysis.

Processing of peptide extracts for liquid chromatography tandem mass spectrometry (LC-MS/MS) analysis included 1) protein extraction and digestion, 2) peptide labelling and 3) peptide fractionation. Peptide labelling was conducted with 8-plex iTRAQ reagents, with a pooled mixture of 4 samples for each tag from I113 to I119. A reference sample with pooled mixture of 112 samples (including additional 28 samples from medicated T2D individuals) was labelled by tag I121 and served as a reference for comparison between the samples from NGT, Pre-DM and TN-T2D individuals. Peptide fractionation was conducted using strong cation-exchange chromatography (SCX).

Bioinformatic analysis: A two-step database search based on the MS/MS spectra was applied, first with a Mascot search against human and gut microbial protein sequence databases to yield a set of scored peptide-spectrum matches (PSMs), followed by a target-decoy search strategy to control for false positives. Identified microbial proteins went through a clustering process using a parsimony principle to generate a non-redundant set of meta-proteins. Both human proteins and meta-proteins were quantified and functionally annotated. Taxonomic assignment was also carried out on meta-proteins.

**Figure S1.** Related to Figure 1: **Concordance and discordance of gut microbiome features in metagenomes and metaproteomes**

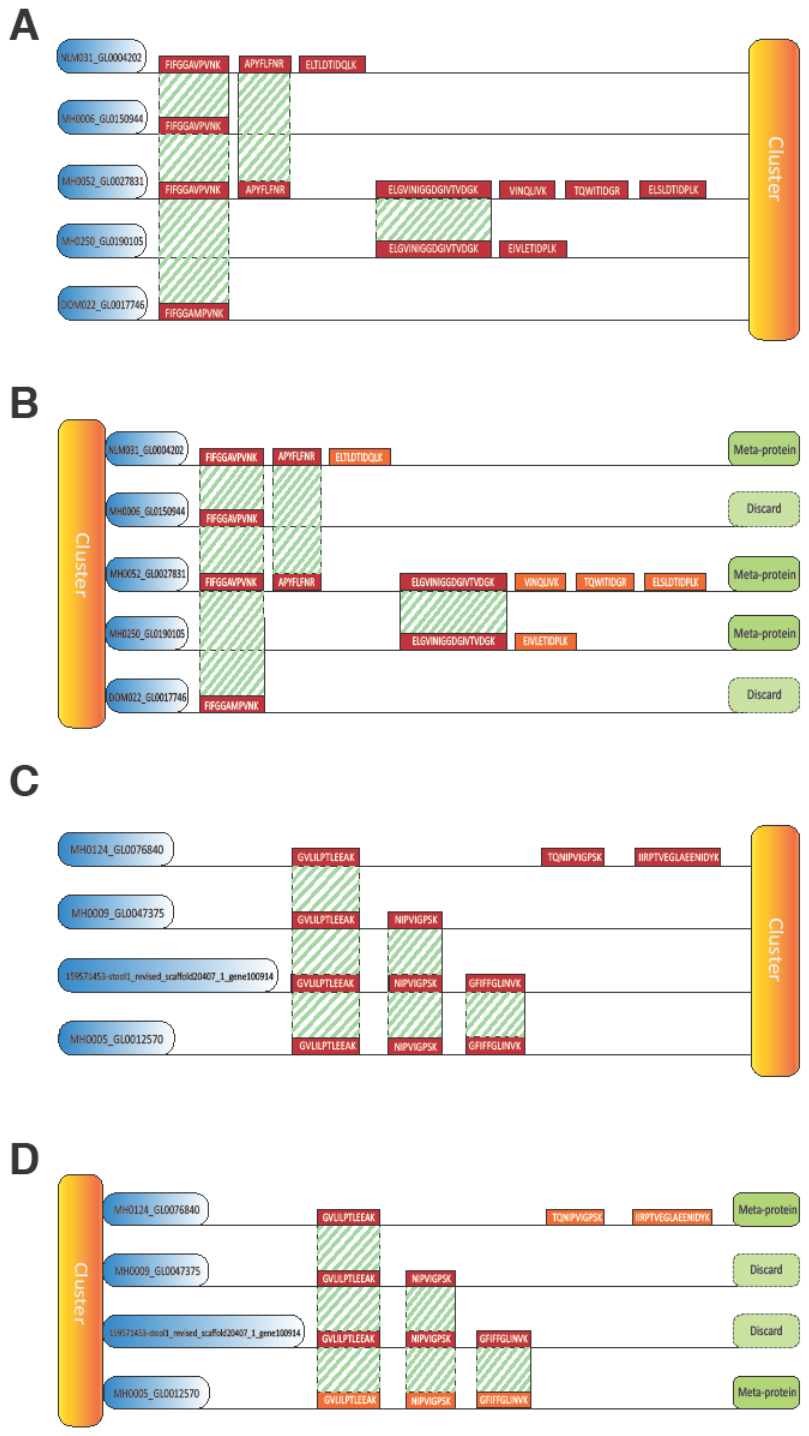

**Figure S2: Meta-protein generation rules**

**(A)** Microbe proteins (left, blue coloured) with at least one shared peptide were grouped into a protein cluster (right, orange coloured). The protein IDs in blue boxes were extracted from potentially encoding genes from the integrated gene catalogue (IGC).

**(B)** The protein cluster identified in (A) was processed according to a maximum parsimony principle, with the minimum number of protein sets containing all the peptides of the cluster retained and defined as meta-protein (right, green coloured). A meta-protein is represented by its encoding gene ID and quantified based on its unique peptides set (coloured in yellow).

**(C-D)** In the case of multiple proteins hits by the same set of peptides and without unique peptides (the last two proteins in panel C and D), the meta-protein was selected as the microbial protein corresponding to the encoding gene with the highest occurrence in the metagenomes.

**Figure S2.** Related to Figure 1: **Experimental overview** and Figure S1.

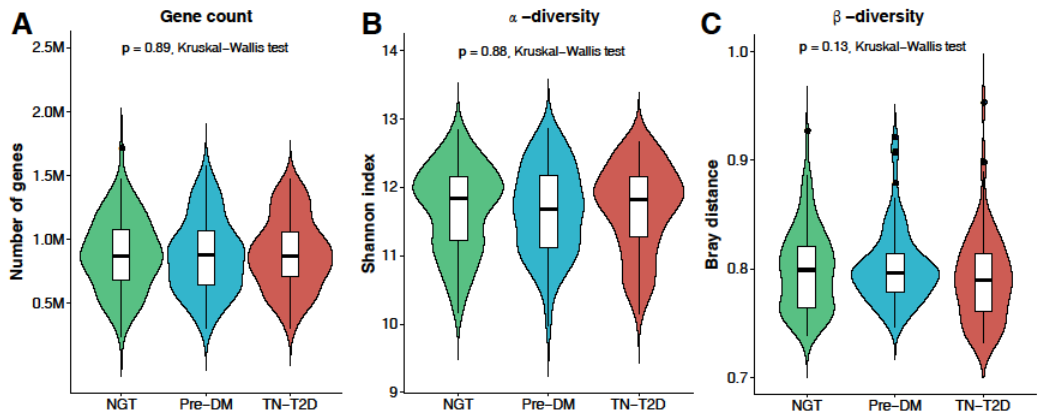

**Figure S3: Summary of richness and diversity measurements**

**(A)** Comparison of gene count in subjects of the three diagnostic groups

**(B)** Comparison of gene-based Shannon indices in subjects of the three diagnostic groups.

**(C)** Comparison of gene-based Bray–Curtis dissimilarity in subjects of the three diagnostic groups.

P values were calculated by Kruskal–Wallis tests.

**Figure S3.** Related to Figure 2: **Determination of alterations in the abundance of MLGs and functional modules**

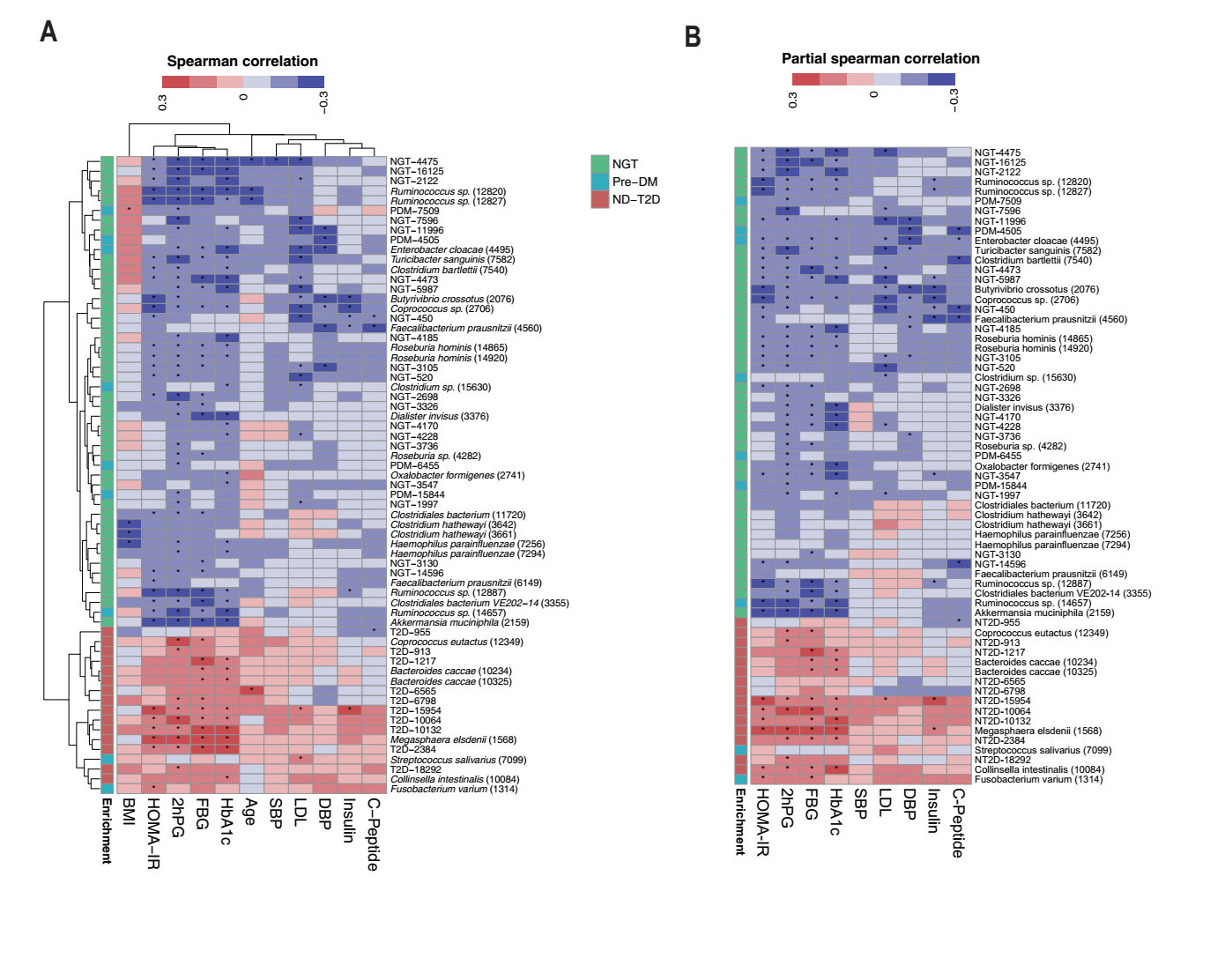

**Figure S4: Correlations between MLGs and phenotypic parameters**

**(A) Spearman’s rank correlation.**

**(B) Partial Spearman’s rank correlation adjusting for age, sex and BMI.**

Taxonomic annotated MLGs displaying significantly differential abundances between NGT, Pre-DM and ND-T2D are shown. Colours represent enrichment in NGT (green), Pre-DM (blue) and ND-T2D (red). *****, correlations with adjusted *P* value < 0.05.

**Figure S4.** Related to Figure 2: **Determination of alterations in the abundance of MLGs and functional modules**

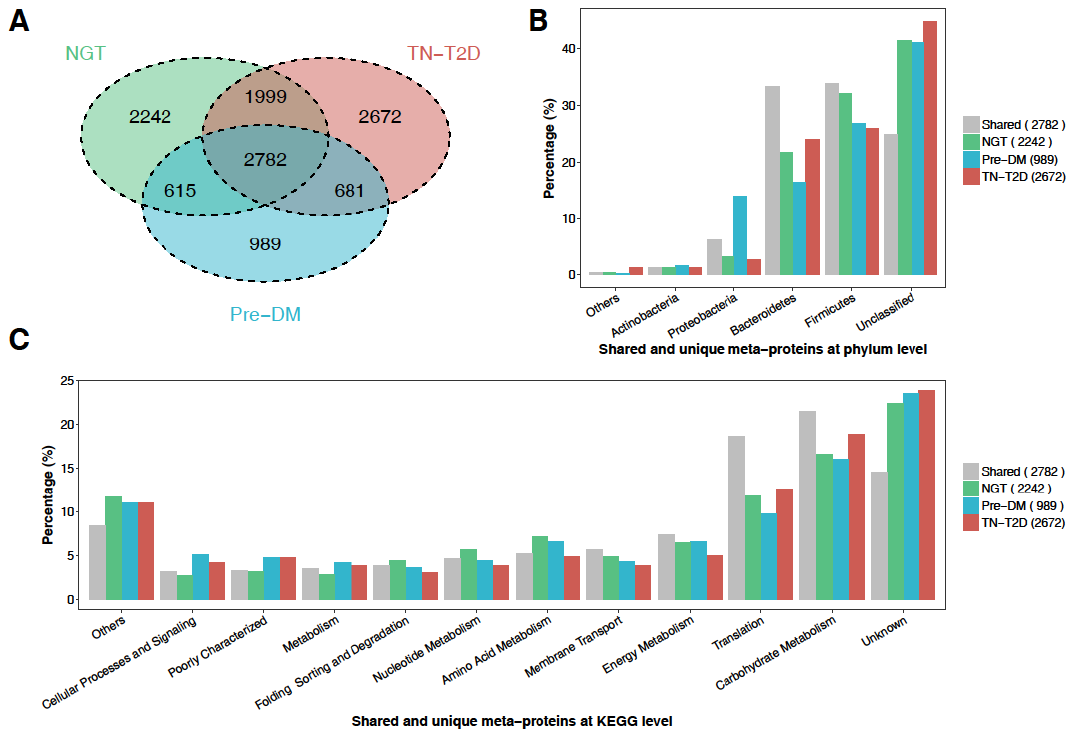

**Figure S5: Distribution of meta-proteins in the three diagnostic groups**

**(A)** Venn diagrams showing the meta-proteins identified in three diagnostic groups.

**(B-C)** The distribution of taxonomic (b) and functional (c) assignment of shared and unique meta-proteins in the three diagnostic groups. A “Others” category indicates other phyla or functional categories accounting for less than 1% of the total microbial community. Colours represent NGT (green), Pre-DM (blue), TN-T2D (red) and shared meta-proteins among the groups (grey).

**Figure S5.** Related to Figure 3A, 3C: **Concordance and discordance of gut microbiome features in metagenomes and metaproteomes**

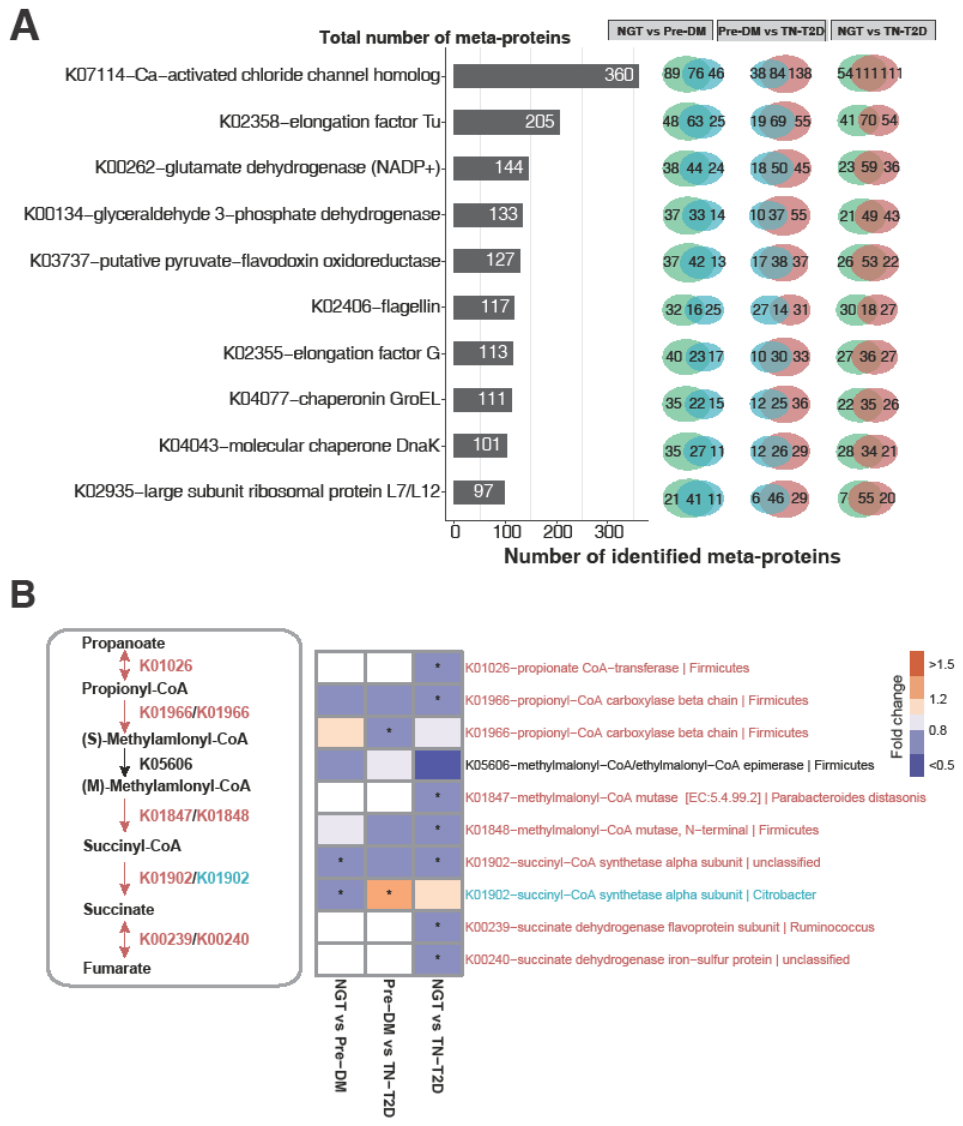

**Figure S6: Identified microbial functions in human faecal samples**

**(A)** Top 10 identified microbial functions in human faecal samples

Bar plot showing the number of meta-proteins in the 10 microbial KOs harbouring the highest number of meta-proteins. Venn diagrams showing the number of KOs identified in three groups and shared between each two groups. Colours represent meta-proteins identified in NGT (green), Pre-DM (blue) and TN-T2D (red) individuals.

**(B)** Abundances of meta-proteins involved in succinate metabolism. Succinate metabolism-related meta-proteins with abundances significantly lower in NGT compared to prediabetes or TN-T2D individuals. Colours represent enrichment in Pre-DM (blue) and TN-T2D (red).

*, *P* < 0.05 and fold change of protein levels > 1.2 or <0.8. Pathways were adapted from KEGG pathways (map00620 and map00650)

**Figure S6.** Related to Figure 3E: **Concordance and discordance of gut microbiome features in metagenomes and metaproteomes**

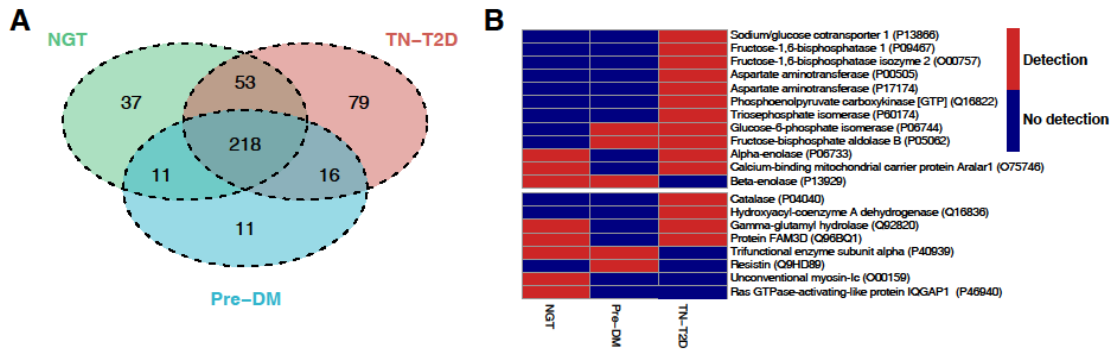

**Figure S7: Distribution of human proteins in the three diagnostic groups**

**(A)** Venn diagram showing the human proteins identified in three diagnostic groups.

**(B)** Detection of human proteins involved in glucose metabolism and insulin signalling pathway.

**Figure S7.** Related to Figure 4: **Characterisation of human proteins in faecal samples from Chinese NGT, Pre-DM, and TN-T2D individuals**

**Supplementary Tables**

**Table S1:** Summary of phenotypic information on the Suzhou T2D cohort

**Table S2:** Metagenomic sequencing statistics for faecal samples from the Suzhou T2D cohort

**Table S3:** List of 84 samples selected for metaproteomics

**Table S4:** Mascot search parameters used in this study

**Table S5:** List of 126 MLGs identified in the Suzhou T2D cohort

**Table S6:** List of published diabetes-associated gut microbial taxa in Chinese and Caucasian

**Table S7:** Accuracy of random forest classifiers in T2D prediction

**Table S8:** Differential enrichment of KEGG modules in individuals with NGT, Pre-DM and TN-T2D

**Table S9:** Summary of proteins, peptides and spectra identified by iTRAQ

**Table S10:** Number of identified meta-proteins assigned to the genus level

**Table S11:** KO assignment of identified meta-proteins

**Table S12:** List of meta-proteins that differed in abundance between individuals with NGT, Pre-DM and TN-T2D

**Table S13:** List of 425 human faecal proteins identified in the Suzhou cohort

**Table S14:** List of human proteins identified in published faecal metaproteomic studies

**Table S15:** List of human proteins that differed in abundance between individuals with NGT, Pre-DM and TN-T2D
